# MinoTour, real-time monitoring and analysis for Nanopore Sequencers

**DOI:** 10.1101/2021.09.10.459783

**Authors:** Rory Munro, Roberto Santos, Alexander Payne, Teri Forey, Solomon Osei, Nadine Holmes, Matt Loose

**Affiliations:** University of Nottingham; Teri Apps: Cork, Co. Cork, IE

**Author notes:** Joint first authors (Rory Munro, Roberto Santos).

## Abstract

**Summary:** MinoTour offers a LIMS system for Oxford Nanopore Technology (ONT) sequencers, with real-time metrics and analysis available permanently for review. Integration of unique real-time automated analysis can reduce the time required to answer biological questions, including mapping and classification of sequence whilst a run is in progress. Real-time sequence data requires new methods of analysis which do not wait for the completion of a run and MinoTour provides a framework to allow users to exploit these features.

**Availability and Implementation:** Source code and documentation are available at https://github.com/LooseLab/minotourcli and https://github.com/LooseLab/minotourapp. Docker images are available from https://hub.docker.com/r/adoni5/, and can be installed using a preconfigured docker-compose script at https://github.com/LooseLab/minotour-docker. An example server is available at http://137.44.59.170.

## Introduction

High throughput real-time portable sequencing has transformed next generation sequencing from specialised centres to individual laboratories and previously unimaginable locations (Quick *et al*., 2016; Castro-Wallace *et al*., 2017). Uniquely, Oxford Nanopore Technologies (ONT) sequencing enables true real-time analysis as sequence data are made available during, as well as after, a run (Loose *et al*., 2016; Payne *et al*., 2020). Rapid analysis of data provides advantages where time to answer is important, such as pathogen genomics and clinical diagnosis of disease (Giovanetti *et al*., 2019; Quick *et al*., 2016; Sone *et al*., 2019; Martignano *et al*., 2021).

During a sequencing run, sequencing is controlled and tracked via ONT’s MinKNOW software. MinKNOW can be used to remotely observe and monitor the progress of sequencing runs, view reports on metrics and recent historical run data. Numerous tools provide analysis of Nanopore FASTQ data either during or after a run (De Coster *et al*., 2018; Bruno *et al*., 2021). ONT also provides a cloud based service, epi2me (EPI2ME), which enables various automated downstream analyses. However, as we show below, minoTour uniquely captures both real-time sequence data and run metrics, can provide analysis and also acts as a comprehensive run archive.

MinoTour is open source and extensible, written using the Django framework, providing real-time insights into sequencer performance and sequence analysis. Visualised metrics allow users to see flow cell performance in real-time. Built in pipelines for alignment and metagenomics allow users to see experimental results in real-time. Additional pipelines can easily be incorporated. For example we can include a customised version of the ARTIC pipeline for SARS-CoV2 analysis described in more detail elsewhere. MinoTour also provides built-in adaptive sequencing (ReadFish only) (Payne *et al*., 2020) support in visualisations enabling users to monitor targeted sequencing in real time.

## Materials and Methods

Figure 1 illustrates the path data takes from the sequencer to the user. A single python tool, minFQ (https://github.com/LooseLab/minotourcli), collects data from two sources. Firstly, run metrics are collected from MinKNOW via an ONT provided application programming interface (API). Secondly base-called data are read either from user specified locations or found via the ONT API. These data are sent to the minoTour server, which saves the sequencer metrics into the database, and stores the sequence data in Redis, a cache database. Celery performs asynchronous analysis of read data and can apply custom pipelines to perform analyses, ranging from base-called data summarisation to sequence alignment, metagenomics or custom workflows such as ARTIC (Tyson *et al*., 2020). Read data are split into individual flow cells, which can be selected from a table to show all data held for individual runs using that flow cell (Figure S1). These data can optionally be shared with other minoTour users if desired (Figure S2). minoTour can also be configured to use twitter/email APIs and notify users about flow cell events such as disk shortages or reaching prespecified coverage thresholds for a reference genome (Figures S3, S4). Reverse communication with the MinKNOW API allows users to send messages back into the MinKNOW logs for a permanent record such as notes on library reloading or comments on flow cell performance.

**Figure 1.**
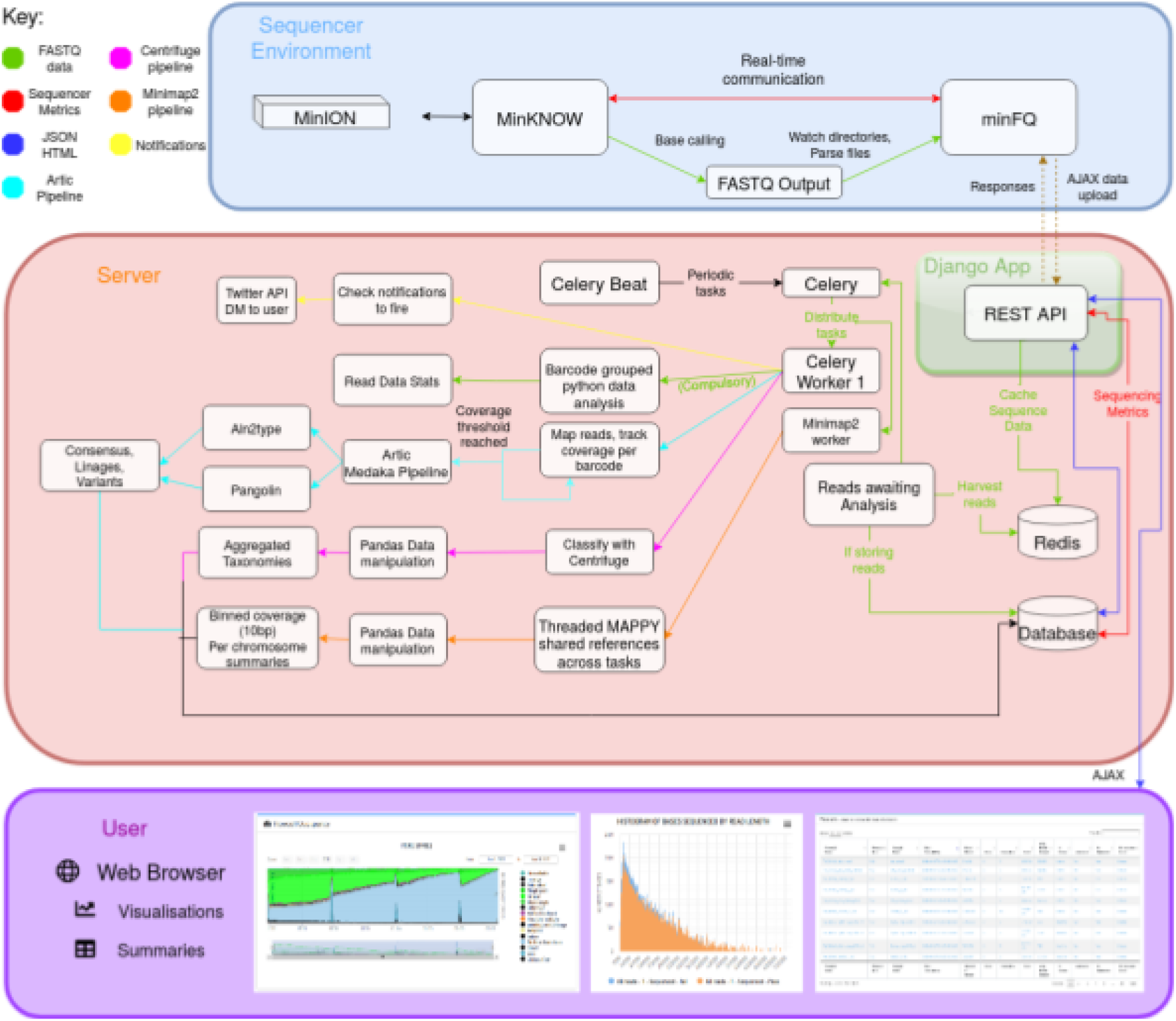
Implementation of minoTour showing the flow of base-called FASTQ data and sequencer metrics from the sequencer to the server, through various optional pipelines and finally to visualisation for the user.

## Results and Discussion

One of minoTour’s strengths is its ability to automatically provide a historic record of runs (Figure S1), and provide real-time key insights into ongoing runs (Figure S5, S6). minoTour also provides a simple overview of all ongoing connected sequencers, quickly showing a user if any are underperforming, such as falling short on yield or speed, who can then respond accordingly (Figure S7). minoTour can also remotely stop a run if required via the website. Using minoTour’s minimap2 alignment pipeline for monitoring targeted or adaptive sequencing with ReadFish one can rapidly tell if coverage is accruing over specific targeted regions with sequenced and rejected reads able to be visualised separately (Figure S8).

minoTour provides a simple metagenomics pipeline, using centrifuge (Kim *et al*., 2016) to visualise sample make-up, a breakdown of broad composition as well as more detailed investigation of select pre-chosen species by aligning reads classified in that taxa against one or more references (Figure S9). More complex pipelines such as the ARTIC pipeline can be incorporated (Figure S10). We routinely use minoTour in the laboratory to monitor sequencing and use the available analyses to provide insights into sequencing experiments. The use of the Django framework enables other users to extend and develop minoTour at will. minFQ, once installed and activated, remains in the background uploading data to the minoTour server, automatically detecting new runs and sequence data (Figure S11).

Our implementation of minoTour with docker alongside detailed server installations for development leave the user with a variety of installation options. MinoTour has been run on large centralised servers, or locally on laptops, enabling monitoring of sequencing runs for collaborative groups, sequencing facilities and individual users. At this time minoTour supports all platforms running flongle or minION flowcells. In future, support for PromethION will become available.

## Acknowledgements

Many thanks to Martin Blythe, Mike Stout and Sunir Malla for their input into this project. We thank Clarie Lonsdale for useful discussions and testing.

## Funding

Work on minoTour has been funded by BBSRC (BB/M020061/1) as well as additional support from the Defence Science and Technology Laboratory (DSTLX-1000138444).

## Conflict of Interest

ML was a member of the MinION access program and has received free flow cells and sequencing reagents in the past. ML has received reimbursement for travel, accommodation and conference fees to speak at events organized by Oxford Nanopore Technologies.

## References

Bruno, A. et al. (2021) BoardION: real-time monitoring of Oxford Nanopore sequencing instruments. BMC Bioinformatics, 22, 245.

Castro-Wallace, S.L. et al. (2017) Nanopore DNA Sequencing and Genome Assembly on the International Space Station. Sci. Rep., 7, 18022.

De Coster, W. et al. (2018) NanoPack: visualizing and processing long-read sequencing data. Bioinformatics, 34, 2666–2669.

EPI2ME https://epi2me.nanoporetech.com/.

Giovanetti, M. et al. (2019) Acute Vector-Borne Viral Infection: Zika and MinION Surveillance. Microbiology spectrum, 7.

Kim, D. et al. (2016) Centrifuge: rapid and sensitive classification of metagenomic sequences. Genome Res., 26, 1721–1729.

Loose, M. et al. (2016) Real-time selective sequencing using nanopore technology. Nat. Methods, 13, 751–754.

Martignano, F. et al. (2021) Nanopore sequencing from liquid biopsy: analysis of copy number variations from cell-free DNA of lung cancer patients. Mol. Cancer, 20, 32.

Payne, A. et al. (2020) Readfish enables targeted nanopore sequencing of gigabase-sized genomes. Nat. Biotechnol., 1–9.

Quick, J. et al. (2016) Real-time, portable genome sequencing for Ebola surveillance. Nature, 530, 228–232.

Sone, J. et al. (2019) Long-read sequencing identifies GGC repeat expansions in NOTCH2NLC associated with neuronal intranuclear inclusion disease. Nat. Genet., 51, 1215–1221.

Tyson, J.R. et al. (2020) Improvements to the ARTIC multiplex PCR method for SARS-CoV-2 genome sequencing using nanopore. bioRxiv.

