## Supplementary Figures for "MinoTour, real-time monitoring and analysis for Nanopore Sequencers"

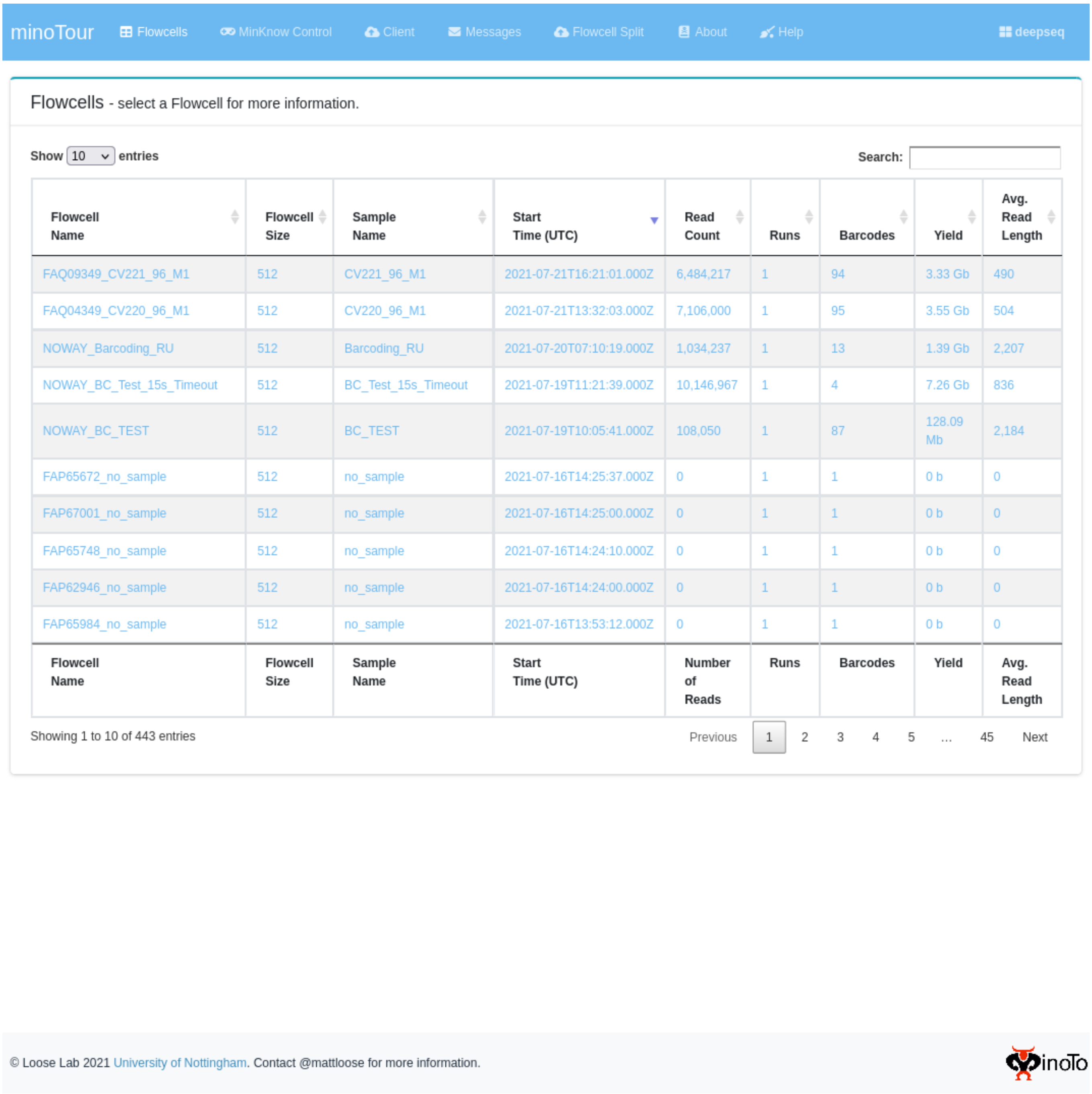


| Flowcell Name | Flowcell Size | Sample Name | Start Time (UTC) | Read Count | Runs | Barcodes | Yield | Avg. Read Length |
|---|---|---|---|---|---|---|---|---|
| FAQ09349_CV221_96_M1 | 512 | CV221_96_M1 | 2021-07-21T16:21:01.000Z | 6,484,217 | 1 | 94 | 3.33 Gb | 490 |
| FAQ04349_CV220_96_M1 | 512 | CV220_96_M1 | 2021-07-21T13:32:03.000Z | 7,106,000 | 1 | 95 | 3.55 Gb | 504 |
| NOWAY_Barcoding_RU | 512 | Barcoding_RU | 2021-07-20T07:10:19.000Z | 1,034,237 | 1 | 13 | 1.39 Gb | 2,207 |
| NOWAY_BC_Test_15s_Timeout | 512 | BC_Test_15s_Timeout | 2021-07-19T11:21:39.000Z | 10,146,967 | 1 | 4 | 7.26 Gb | 836 |
| NOWAY_BC_TEST | 512 | BC_TEST | 2021-07-19T10:05:41.000Z | 108,050 | 1 | 87 | 128.09 Mb | 2,184 |
| FAP65672_no_sample | 512 | no_sample | 2021-07-16T14:25:37.000Z | 0 | 1 | 1 | 0 b | 0 |
| FAP67001_no_sample | 512 | no_sample | 2021-07-16T14:25:00.000Z | 0 | 1 | 1 | 0 b | 0 |
| FAP65748_no_sample | 512 | no_sample | 2021-07-16T14:24:10.000Z | 0 | 1 | 1 | 0 b | 0 |
| FAP62946_no_sample | 512 | no_sample | 2021-07-16T14:24:00.000Z | 0 | 1 | 1 | 0 b | 0 |
| FAP65984_no_sample | 512 | no_sample | 2021-07-16T13:53:12.000Z | 0 | 1 | 1 | 0 b | 0 |
| Flowcell Name | Flowcell Size | Sample Name | Start Time (UTC) | Number of Reads | Runs | Barcodes | Yield | Avg. Read Length |



**Supplementary figure 1 - Flow cell table.** The landing page for an authenticated user, displaying quick metrics about flow cells.The table is sortable and searchable. Upon clicking a row, the user is redirected to a tabbed page displaying all data held about the flow cell.

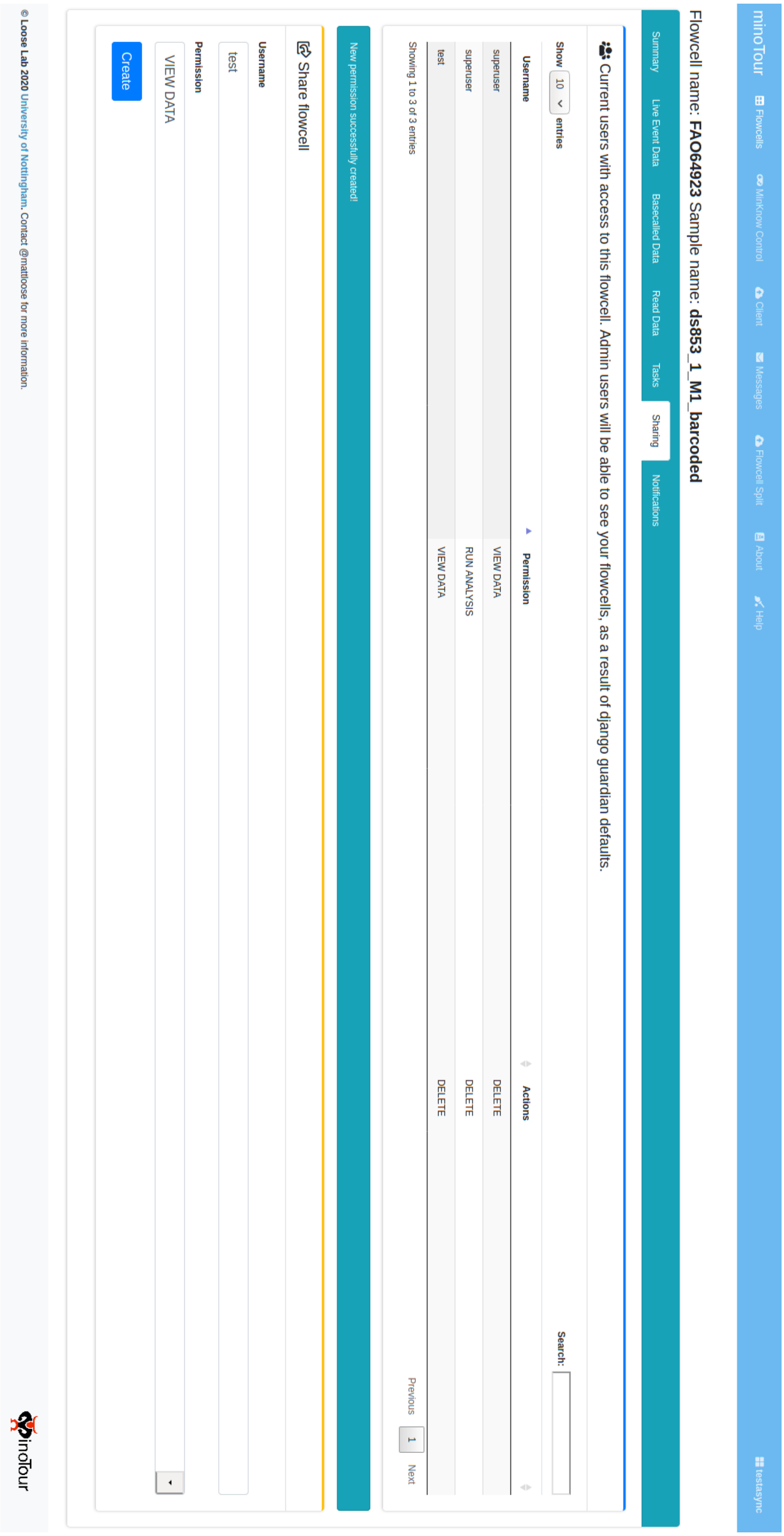


**Supplementary figure 2 - Sharing tab**. Uploaded flow cells can be shared with other users, with a permission level of either VIEW DATA, or RUN ANALYSES. The sharing permission can later be deleted. Admin users can see all flow cells that have been shared, due to an unalterable default in the package used to manage sharing permissions.

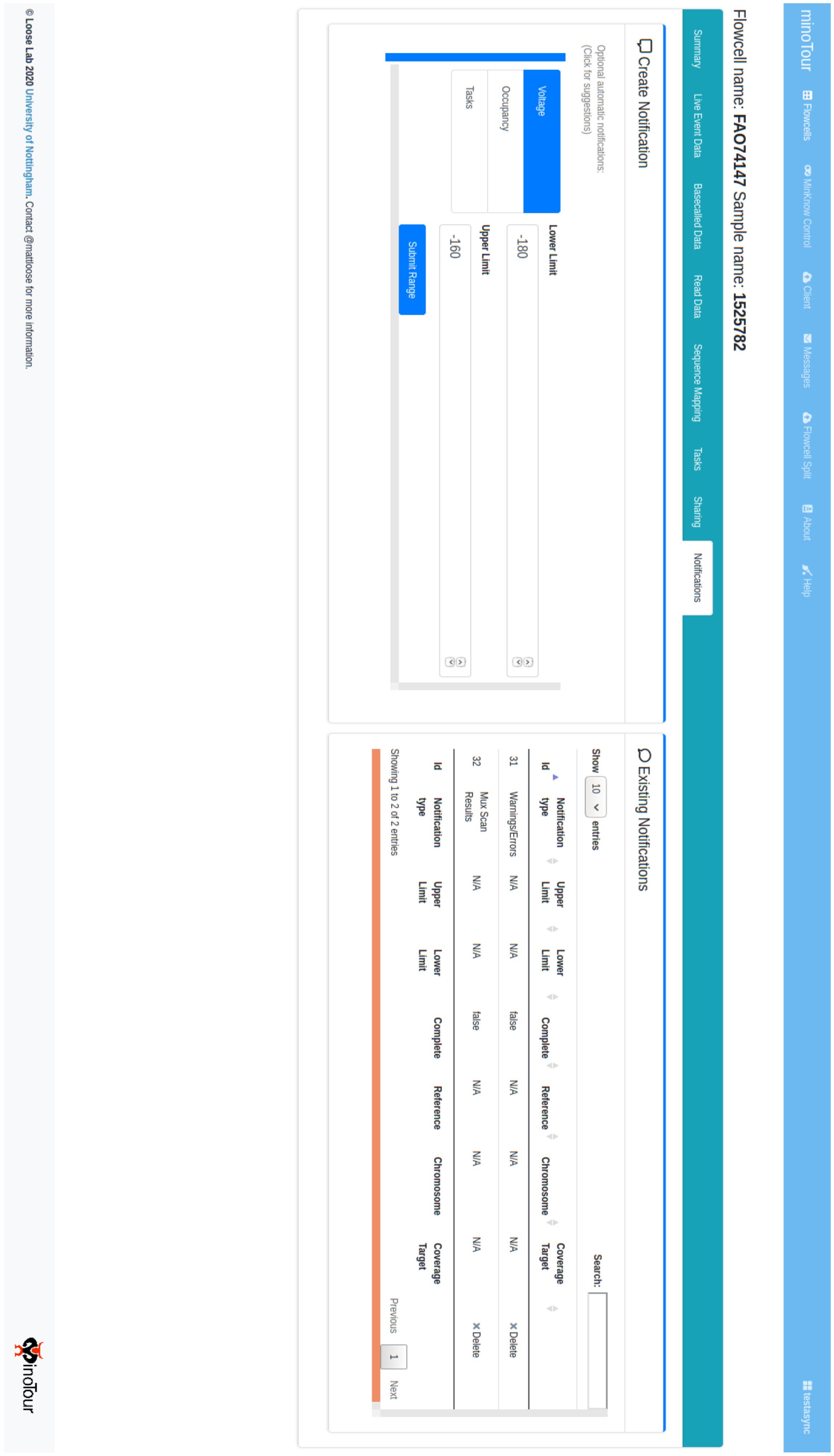


**Supplementary figure 3 - Notifications tab**. Users can enable notifications that are triggered when certain conditions are met. For example when MinKNOW performs a mux scan, the results of this mux scan are sent to the user via Twitter, if the user has provided their twitter details.

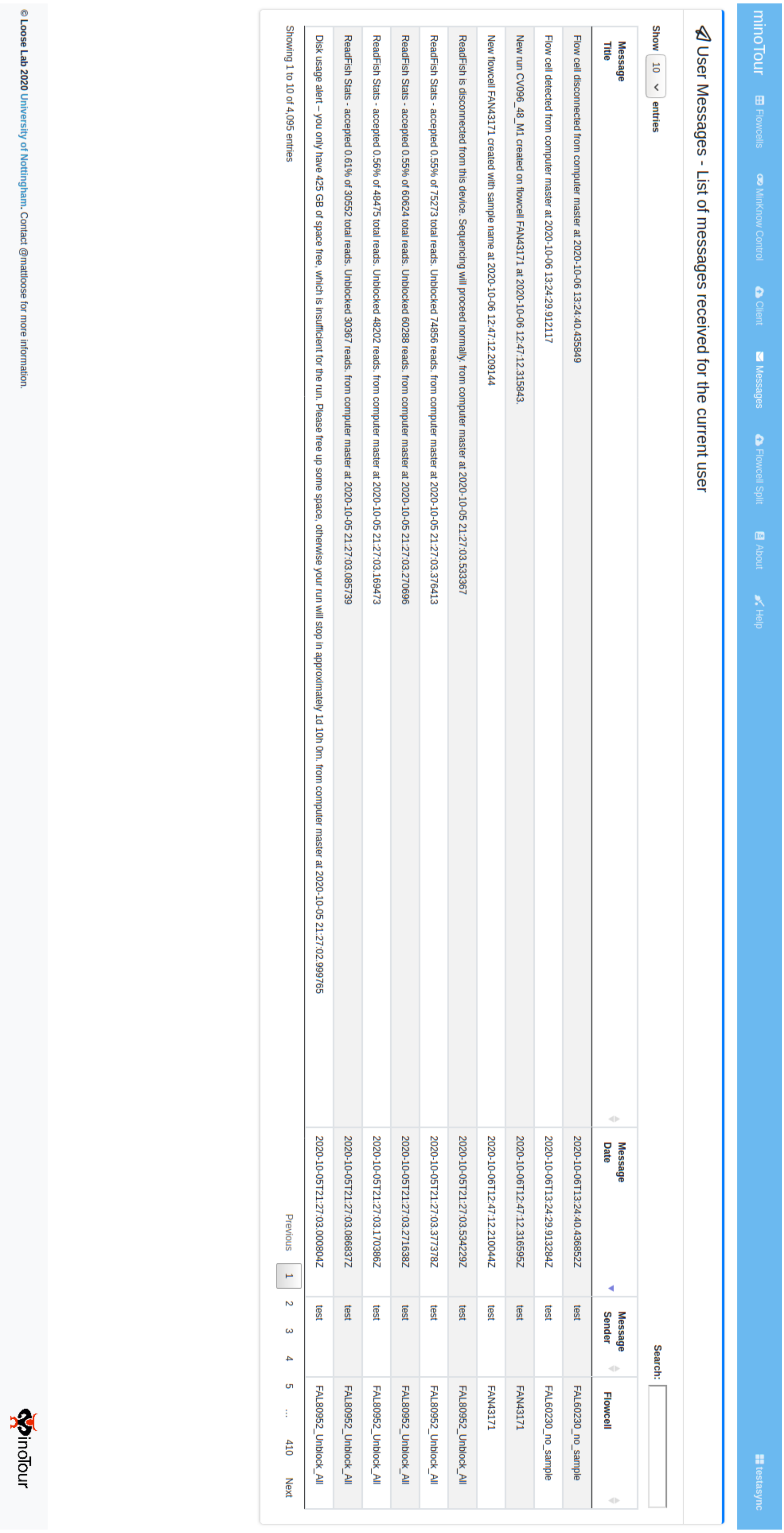


| Message Title | Message Date | Message Sender | Flowcell |
|---|---|---|---|
| Flow cell disconnected from computer master at 2020-10-06 13:24:40.435849 | 2020-10-06T13:24:40.436852Z | test | FAL60230_no_sample |
| Flow cell detected from computer master at 2020-10-06 13:24:29.912117 | 2020-10-06T13:24:29.913284Z | test | FAL60230_no_sample |
| New run CV096_48_M1 created on flowcell FAN43171 at 2020-10-06 12:47:12.315843. | 2020-10-06T12:47:12.316595Z | test | FAN43171 |
| New flowcell FAN43171 created with sample name at 2020-10-06 12:47:12.209144 | 2020-10-06T12:47:12.210044Z | test | FAN43171 |
| ReadFish is disconnected from this device. Sequencing will proceed normally. from computer master at 2020-10-05 21:27:03.533367 | 2020-10-05T21:27:03.534229Z | test | FAL80952_Unblock_All |
| ReadFish Stats - accepted 0.55% of 75273 total reads. Unblocked 74856 reads. from computer master at 2020-10-05 21:27:03.376413 | 2020-10-05T21:27:03.377378Z | test | FAL80952_Unblock_All |
| ReadFish Stats - accepted 0.55% of 60624 total reads. Unblocked 60288 reads. from computer master at 2020-10-05 21:27:03.270696 | 2020-10-05T21:27:03.271638Z | test | FAL80952_Unblock_All |
| ReadFish Stats - accepted 0.56% of 48475 total reads. Unblocked 48202 reads. from computer master at 2020-10-05 21:27:03.169473 | 2020-10-05T21:27:03.170386Z | test | FAL80952_Unblock_All |
| ReadFish Stats - accepted 0.61% of 30552 total reads. Unblocked 30367 reads. from computer master at 2020-10-05 21:27:03.085739 | 2020-10-05T21:27:03.086837Z | test | FAL80952_Unblock_All |
| Disk usage alert – you only have 425 GB of space free, which is insufficient for the run. Please free up some space, otherwise your run will stop in approximately 1d 10h 0m. from computer master at 2020-10-05 21:27:02.999765 | 2020-10-05T21:27:03.000804Z | test | FAL80952_Unblock_All |



**Supplementary figure 4 - Messages page**. When notification conditions are met, or when a new flow cell or sequencing run is created in minoTour, a message is created inside the system. This message is then tweeted to the user, alerting them to certain events in the system.

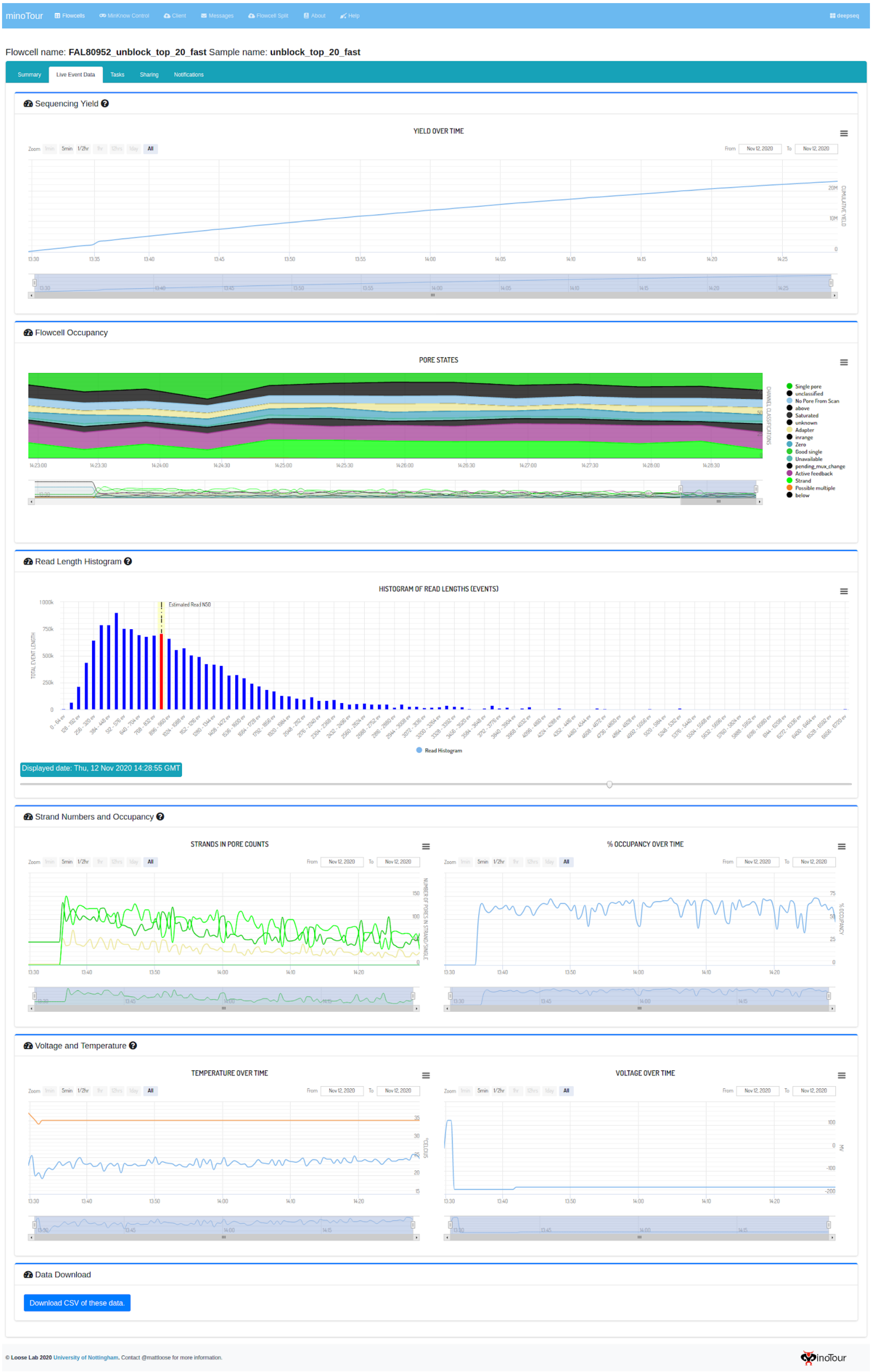


**Supplementary figure 5 - Live sequencing metrics tab**. This page visualises several sequencing metrics historically, allowing a perfect record of sequencing runs to be kept.

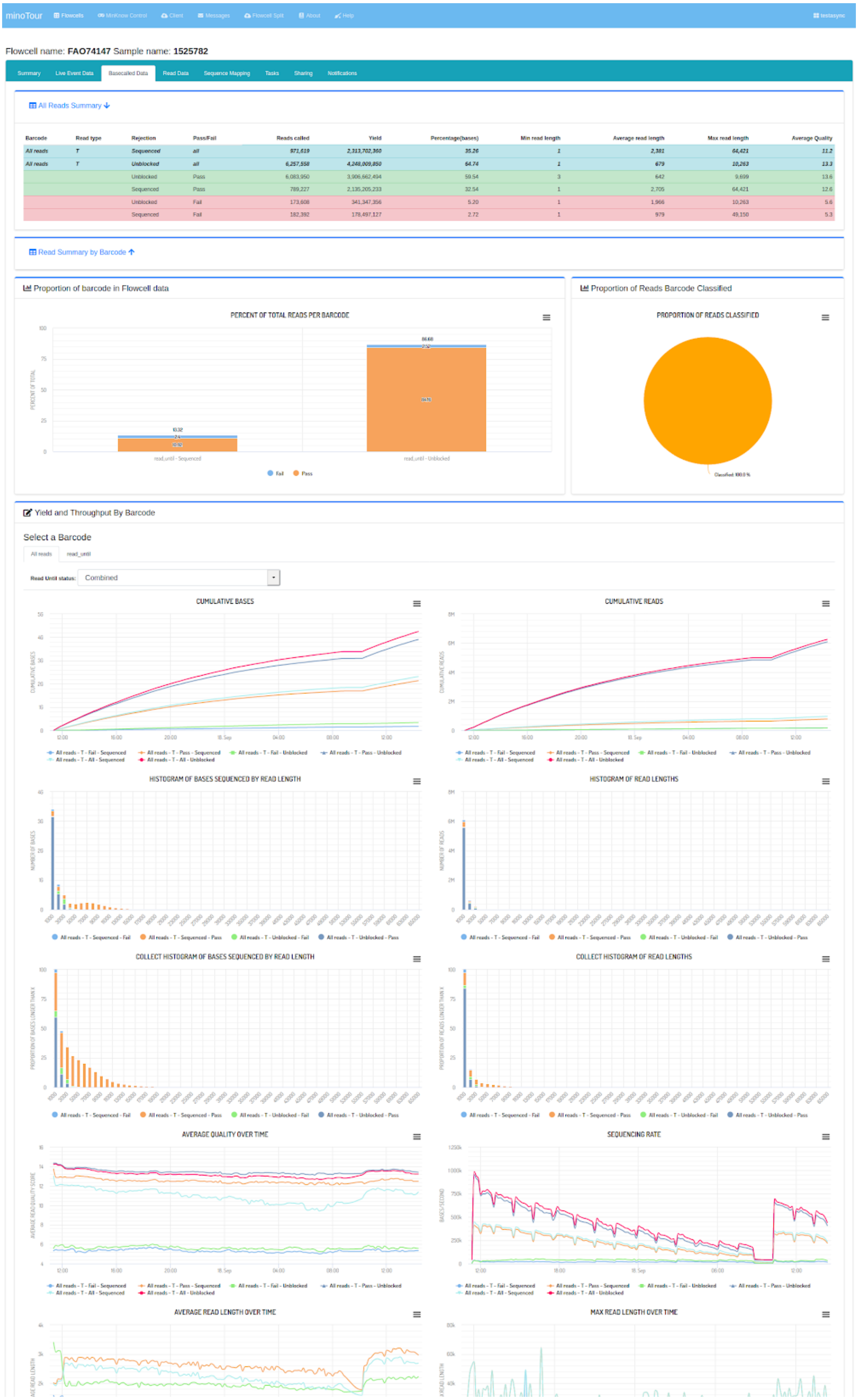


**Supplementary figure 6 - Base-called data tab**. A summary of base called data produced by a sequencing run. This data can be uploaded separately or not at all. Omitted are a tabular summary of data for each barcode and heatmaps of the flow cell showing yield and read count per pore.

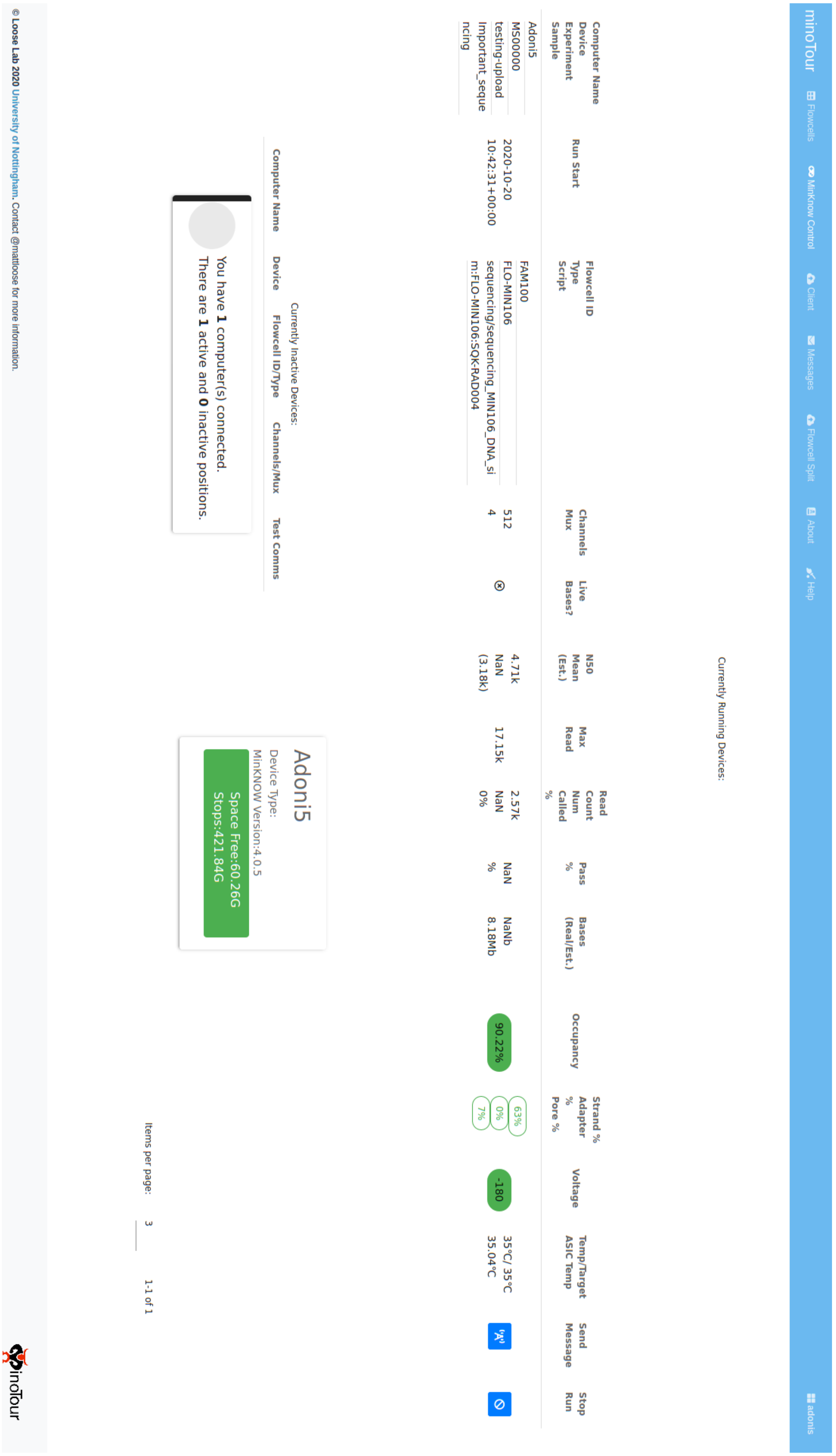


**Supplementary figure 7 - MinKNOW control**. MinoTour offers comprehensive LIMS capabilities. Demonstrated here is the ability to monitor the live performance of all connected devices, easily flagging any issues, send custom messages to the MinKNOW instance that the device is connected to and remotely stop a run.

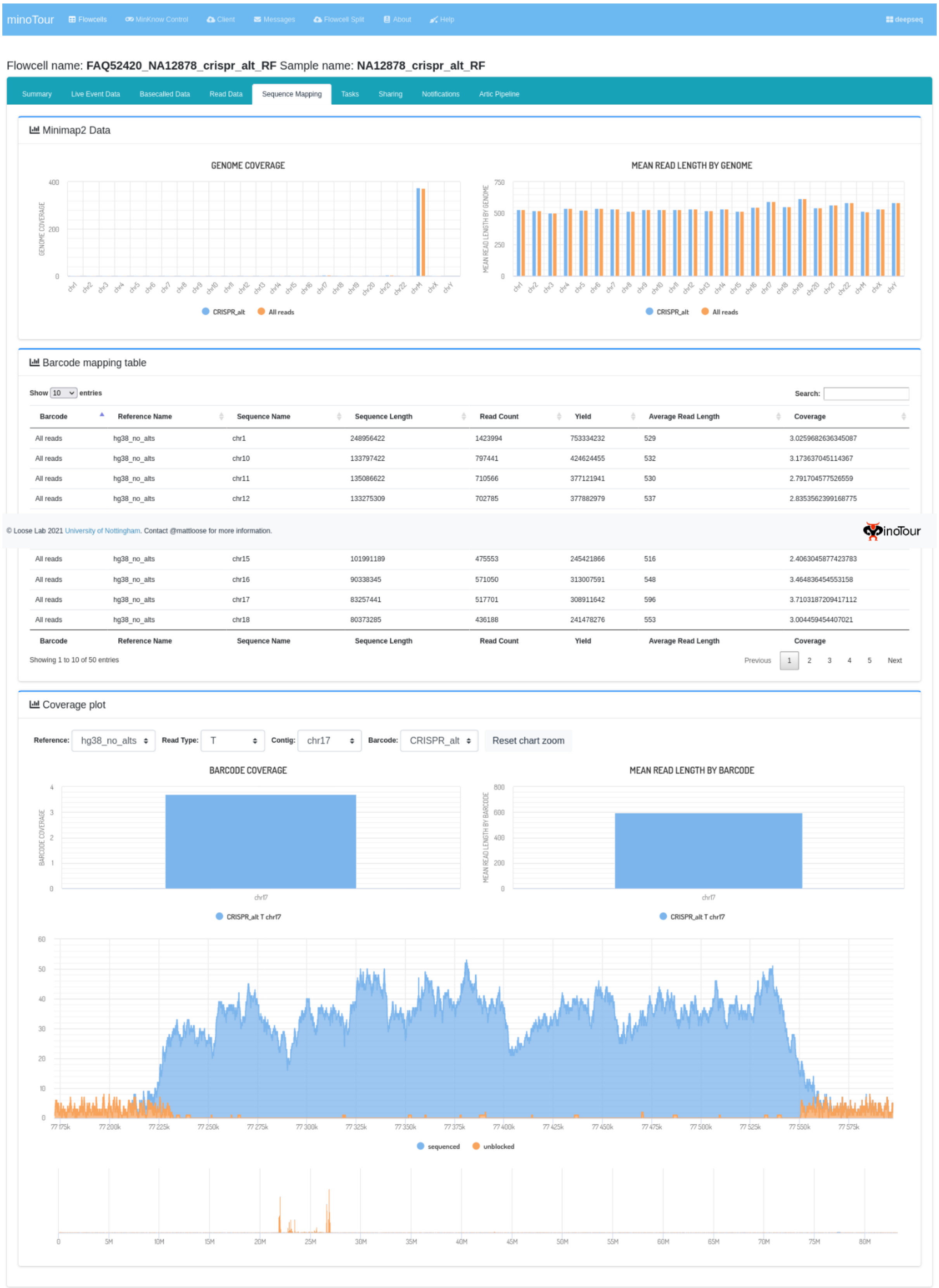


**Supplementary figure 8 - Alignment tab**. The alignment analysis in minoTour offers a quick insight into a sequencing run in real time, aligning to user specified references. The alignments are broken down by barcode and contig (if multiple). This is not intended as a final result, it is merely to flag any areas that would be interesting for further study. In adaptive sequencing, unblocked and sequenced reads are separately visualised.

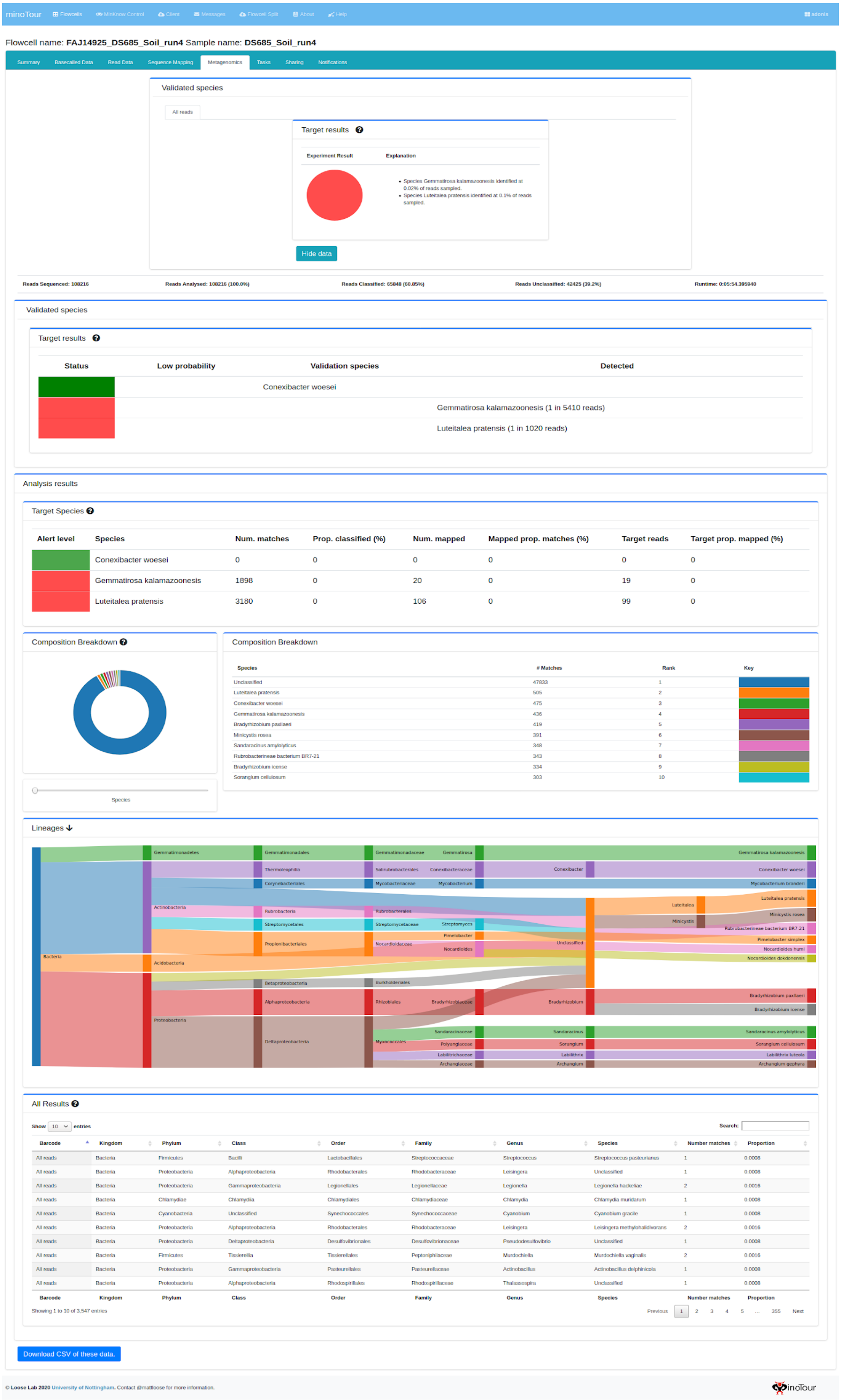


**Supplementary figure 9 - Metagenomics tab.** The metagenomics analysis is a quick insight into the broader composition of a dataset. Visualisations include a Sankey Diagram denoting the lineages of the 30 most common species by read number, a composition breakdown at each taxonomic level and full results table that is searchable and sortable.

minoTour | Flowcells | MinKnow Control | Client | Messages | Flowcell Split | About | Help | adonis

Flowcell name: **FAQ01023_CV197_25_M1** Sample name: **CV197_25_M1**

Summary | Basecalled Data | Read Data | Tasks | Sharing | Notifications | Artic Pipeline

### Artic Coverage Charts

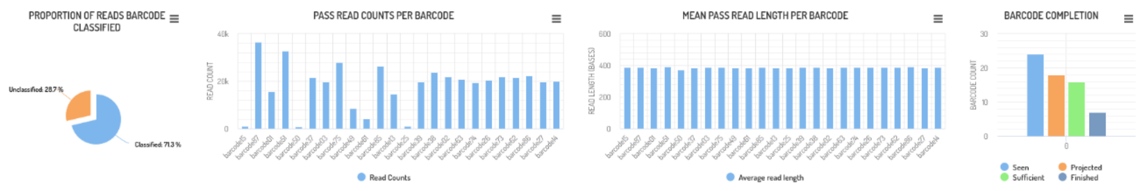


### Artic Phylogeny Reports

A SNIPIT plot (see https://github.com/aineniamh/snipit) for consensus sequences for samples completed so far.
Sequences will be added or updated as consensus sequences are generated over time. When a new sequence is generated, the most complete will be used in this plot.
**Only sequences with less than 50% N calls are included in this analysis.**

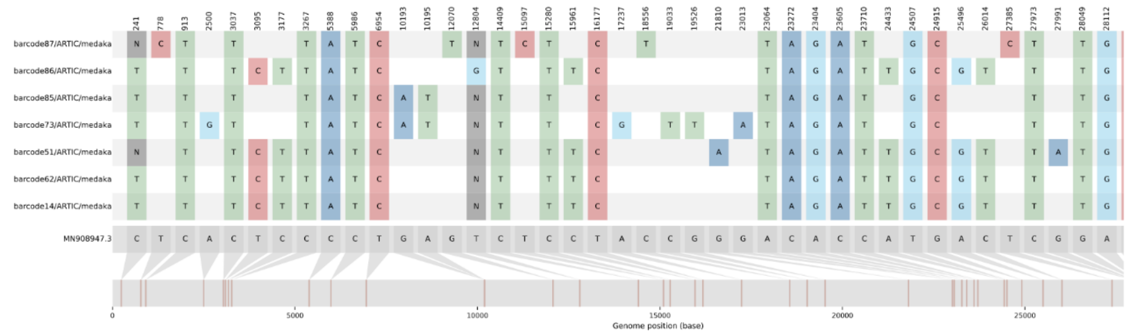


A simple phylogenetic tree for sequences. Trees are generated using iqtree after alignment with mafft. The reference sequence provided for the artic pipeline is used as an outgroup. Trees are visualised using figtree.js.
**Only sequences with less than 50% N calls are included in this analysis. Use the slider to adjust the height of the tree image.**

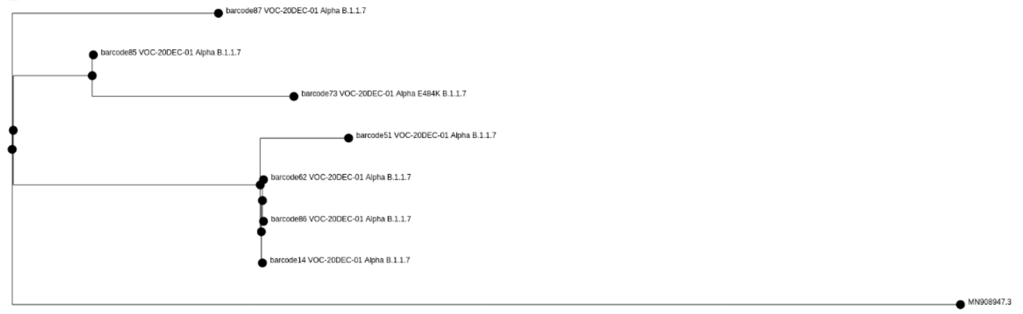


### Pipeline firing conditions

20 X Coverage at 90 % of amplicons Submit choices

Show 10 entries | Search:

| % of amplicons | X coverage | Delete |
|---|---|---|
| 90 | 20 | ✕ Delete |
| % of amplicons | X coverage | Delete |

Showing 1 to 1 of 1 entries | Previous 1 Next

### Summary Table (Pass reads only)

Copy | CSV | PDF | Show 10 rows | Search: barcode

| Barcode | Sequence Name | Sequence Length | Read Count | Yield | Average Read Length | Coverage | # Success Amplicons | # Partial Amplicons | # Failed Amplicons | Mean of amplicons | Var of amplicons | Amplicon Std. Dev | Sufficient Coverage | Projected to Finish | Lineage | VoC Found |
|---|---|---|---|---|---|---|---|---|---|---|---|---|---|---|---|---|
| barcode15 | MN908947.3 | 29,903 | 1,137 | 440.68 kb | 388 | 14.45 | 16 | 4 | 79 | 13.4 | 1102.03 | 33.2 | false | false | Currently unknown | Not Tested |
| barcode87 | MN908947.3 | 29,903 | 36,440 | 14.12 Mb | 388 | 463.22 | 99 | 0 | 0 | 445.87 | 29669.97 | 172.25 | true | true | B.1.1.7 | VOC-20DEC-01 (confirmed) |
| barcode01 | MN908947.3 | 29,903 | 15,766 | 6.09 Mb | 386 | 199.52 | 71 | 9 | 19 | 188.82 | 60892.39 | 246.76 | false | false | Currently unknown | Not Tested |
| barcode51 | MN908947.3 | 29,903 | 32,764 | 12.81 Mb | 391 | 420.15 | 99 | 0 | 0 | 404.61 | 26610.79 | 163.13 | true | true | B.1.1.7 | VOC-20DEC-01 (confirmed) |
| barcode50 | MN908947.3 | 29,903 | 705 | 262.70 kb | 373 | 8.61 | 10 | 9 | 80 | 8.46 | 887.69 | 29.79 | false | false | Currently unknown | Not Tested |
| barcode37 | MN908947.3 | 29,903 | 21,563 | 8.31 Mb | 386 | 272.39 | 89 | 6 | 4 | 261.48 | 46975.64 | 216.74 | false | true | Currently unknown | Not Tested |
| barcode03 | MN908947.3 | 29,903 | 19,712 | 7.63 Mb | 387 | 249.89 | 92 | 7 | 0 | 243.16 | 24450.24 | 156.37 | true | true | Currently unknown | Not Tested |
| barcode75 | MN908947.3 | 29,903 | 28,058 | 10.87 Mb | 388 | 356.25 | 95 | 4 | 0 | 342.77 | 39780.85 | 199.45 | true | true | Currently unknown | Not Tested |
| barcode49 | MN908947.3 | 29,903 | 8,602 | 3.31 Mb | 384 | 108.39 | 56 | 13 | 30 | 108.21 | 22371.43 | 149.57 | false | false | Currently unknown | Not Tested |

| Barcode | Sequence Name | Sequence Length | Read Count | Yield | Average Read Length | Coverage | # Success Amplicons | # Partial Amplicons | # Failed Amplicons | Mean of amplicons | Var of amplicons | Amplicon Std. Dev | Sufficient Coverage | Projected to Finish | Lineage | VoC Found |
|---|---|---|---|---|---|---|---|---|---|---|---|---|---|---|---|---|
| barcode61 | MN908947.3 | 29,903 | 4,319 | 1.67 Mb | 386 | 54.75 | 32 | 6 | 61 | 49.38 | 8000.96 | 89.45 | false | false | Currently unknown | Not Tested |

Showing 1 to 10 of 24 entries (filtered from 25 total entries)

Previous 1 2 3 Next

Download all completed | Run all Incomplete | Re-Run All | Export report

### Lineage Information for barcode87/ARTIC/medaka - B.1.1.7

Links to information on lineage B.1.1.7

PANGO Info | Outbreak Info

### Variant of Concern Report for barcode87/ARTIC/medaka

If a genome is analysed, this will provide a report for the variant or variants found based on current PHE Variant Definitions. This analysis will report all possible VoCs and caution should be taken in interpretation of low coverage/quality genomes.

Reported VuI/VoCs:

| sample_id | phe-label | unique-id | status | mutation-ref-calls | mutation-mixed-calls | mutation-calls | indel-ref-calls | indel-calls | no-calls | no-calls-deletion |
|---|---|---|---|---|---|---|---|---|---|---|
| barcode87/ARTIC/medaka | VOC-20DEC-01 | denture-daughter | confirmed | 0 | 0 | 13 | 0 | 2 | 0 | 0 |

Observed mutations:

| type | snp | snp | snp | snp | snp | snp | snp | del | snp | snp | snp | snp | snp | snp | del |
|---|---|---|---|---|---|---|---|---|---|---|---|---|---|---|---|
| reference-base | T | C | C | C | C | C | T | GTCTGGTTTT | G | C | T | C | T | C | ATACATG |
| variant-base | C | T | T | T | A | T | C | G | T | T | C | T | C | T | A |
| var-length | 1 | 1 | 1 | 1 | 1 | 1 | 1 | 10 | 1 | 1 | 1 | 1 | 1 | 1 | 7 |
| one-based-reference-position | 778 | 913 | 3037 | 3267 | 5388 | 5986 | 6954 | 11287 | 12070 | 14408 | 15096 | 15279 | 16176 | 18555 | 21764 |
| iupac-variant-bases | C | T | T | T | A | T | C | G | T | T | C | T | C | T | A |

| type | del | snp | snp | snp | snp | snp | snp | snp | snp | snp | snp | snp | del | mnp | mnp-snp | mnp-snp |
|---|---|---|---|---|---|---|---|---|---|---|---|---|---|---|---|---|
| reference-base | TTTA | A | C | A | C | C | T | G | T | C | G | A | TA | GAT | A | T |
| variant-base | T | T | A | G | A | T | G | C | C | T | T | G | T | CTA | T | A |
| var-length | 4 | 1 | 1 | 1 | 1 | 1 | 1 | 1 | 1 | 1 | 1 | 1 | 2 | 3 | 1 | 1 |
| one-based-reference-position | 21990 | 23063 | 23271 | 23403 | 23604 | 23709 | 24506 | 24914 | 27384 | 27972 | 28048 | 28111 | 28270 | 28280 | 28281 | 28282 |
| iupac-variant-bases | T | T | A | G | A | T | G | C | C | T | T | G | T | CTA | T | A |

Sample_ID: barcode87/ARTIC/medaka
PANGO:B.1.1.7
nextstrain:N501Y.V1

Variant Status: confirmed

PHE-Label: VOC-20DEC-01
WHO Label: Alpha
Alternate Names: VOC202012/01, UK variant, Kent variant, VOC1,

*Description:*
This variant became widespread in the UK in the Winter of 2021 and is characterised by increased transmissibility.
*Information Sources* Source_1 Source_2

Variant Calls Detected:

| Position | gene | protein | ref | variant | sample call | type | status |
|---|---|---|---|---|---|---|---|
| 3267 | ORF1ab | nsp3 | C | T | T | SNP | detect |
| 5388 | ORF1ab | nsp3 | C | A | A | SNP | detect |
| 6954 | ORF1ab | nsp3 | T | C | C | SNP | detect |
| 21764 | S | surface glycoprotein | ATACATG | A | A | deletion | detect |
| 21990 | S | surface glycoprotein | TTTA | T | T | deletion | detect |
| 23063 | S | surface glycoprotein | A | T | T | SNP | detect |
| 23271 | S | surface glycoprotein | C | A | A | SNP | detect |
| 23604 | S | surface glycoprotein | C | A | A | SNP | detect |
| 23709 | S | surface glycoprotein | C | T | T | SNP | detect |
| 24506 | S | surface glycoprotein | T | G | G | SNP | detect |
| 24914 | S | surface glycoprotein | G | C | C | SNP | detect |
| 27972 | ORF8 | ORF8 protein | C | T | T | SNP | detect |
| 28048 | ORF8 | ORF8 protein | G | T | T | SNP | detect |
| 28111 | ORF8 | ORF8 protein | A | G | G | SNP | detect |
| 28280 | N | nucleocapsid phosphoprotein | GAT | CTA | CTA | MNP | detect |

One based position - notes on sites detected in this sample.

The observed calls were:

- mutation_ref_calls: 0
- indel_ref_calls: 0
- mutation_calls: 13
- mutation_mixed_calls: 0
- indel_calls: 2
- no_calls: 0
- no_call_deletion: 0

Classification rules:

| confirmed | probable | low_qc |
|---|---|---|
| mutations_required: 13 | mutations_required: 5 | mutations_required: 0 |
| indels_required: 0 | indels_required: 0 | indels_required: 0 |
| allowed_wildtype: 0 | allowed_wildtype: 0 | allowed_wildtype: 0 |

*Acknowledgements:*
*Curators:*Natalie Groves Ulf Schaefer Nick Loman

The complete list of variants of concern scanned included:
VUI-21FEB-04 VOC-20DEC-02 VUI-21FEB-01 VUI-21MAR-01 VUI-21MAY-01 VOC-20DEC-01 VOC-21APR-02 VUI-21FEB-03 VUI-21APR-01 E484K VUI-21JUL-01 VUI-21JAN-01 VUI-21MAR-02 VUI-21MAY-02 VUI-21JUN-01 VOC-21JAN-02 VUI-21APR-03 VOC-21FEB-02

### Coverage Plots (Passed Reads)

barcode15 Log Coverage
Select a barcode from the select box and then highlight a region on this coverage chart to see it in detail below.

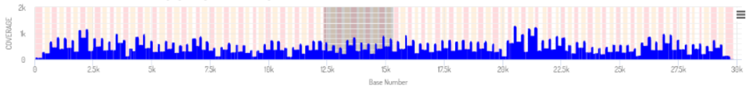


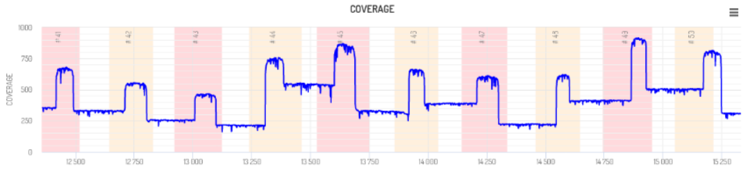


### barcode87 Results Summary

Summary of barcode data.

| | |
|---|---|
| **Avg. Coverage** | 438 |
| **Var. Coverage** | 48615.31 |
| **Min. Coverage** | 0 |
| **Max. Coverage** | 1299 |
| **% reads in run** | 3.42 |
| **Has Finished** | True |
| **Has Sufficient Coverage** | True |

Download Results Files | Re-run Artic Command

Pangolin Lineage Assignments.

| | 0 |
|---|---|
| **taxon** | barcode87/ARTIC/medaka_MN908947.3 |
| **lineage** | B.1.1.7 |
| **conflict** | 0.0 |

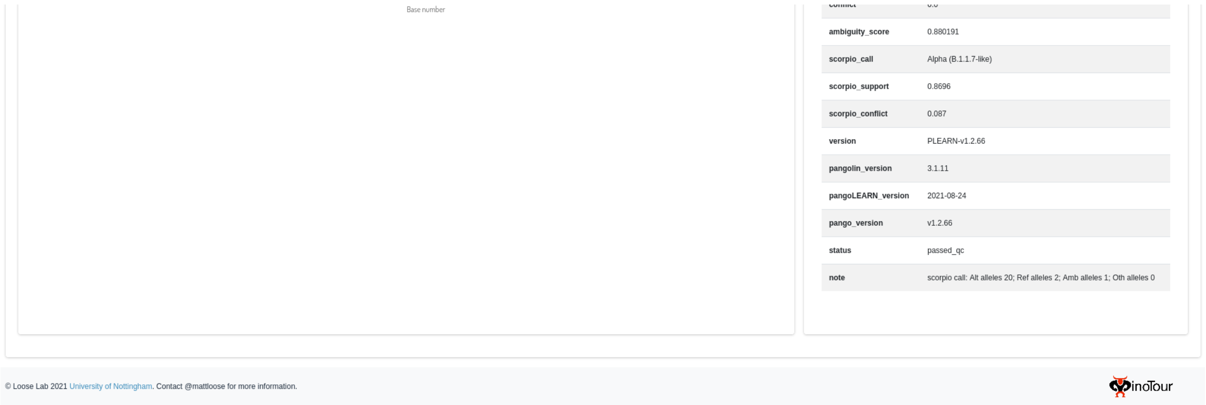


**Supplementary figure 10 - ARTIC Pipeline results.** This page visualises the output of the ARTIC pipeline built into minoTour, showing metrics per sample, lineage for completed samples and variants and a simple phylogenetic tree for all lineages.

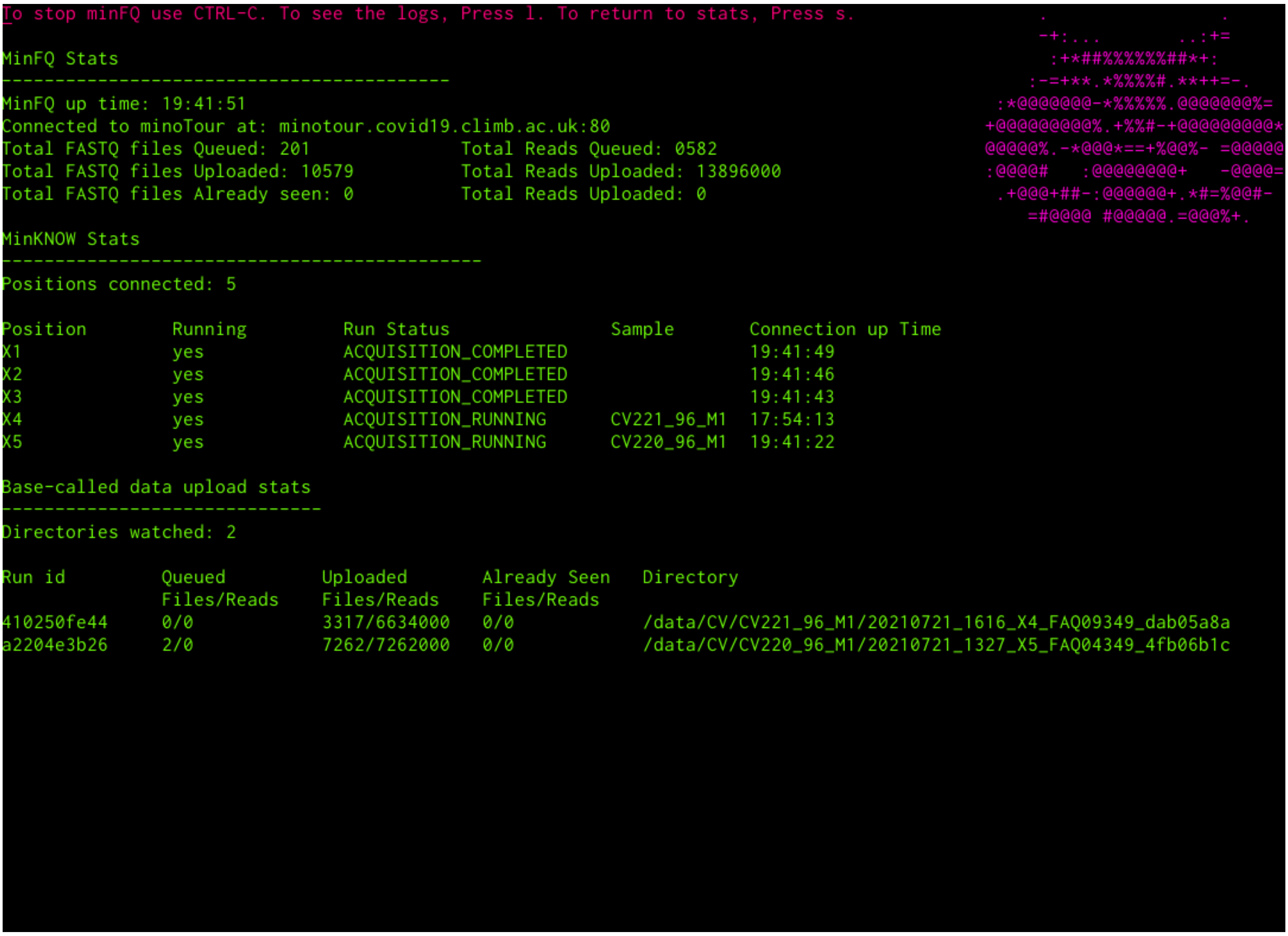


**Supplementary figure 11 - MinFQ terminal interface.** The minFQ interface here lists upload statistics for live monitoring and for base-called data automatically detected from the run.
